# POPULATION CHARACTERISATION AND PROPAGATION STRATEGY FOR WILD *Coffea eugenioides* IN NYUNGWE AND CYAMUDONGO FORESTS, RWANDA: *A field assessment of mother trees and an evidence-based seed-source pathway*

**DOI:** 10.64898/2026.06.16.732672

**Authors:** Yves Shema

## Abstract

This report presents a field assessment of wild *Coffea eugenioides*, the diploid maternal progenitor of cultivated Arabica coffee, in Nyungwe Forest National Park and the adjoining, isolated Cyamudongo forest fragment in south-western Rwanda. The assessment targeted trees identified as candidate mother trees (individuals bearing fruit and therefore potential seed sources): 59 such trees were georeferenced and measured for structural, phenological and reproductive traits using a structured KoBoToolbox questionnaire (reproduced in Annex 1). The work was undertaken to inform the establishment of a seed orchard for this species under the RTRP-Seed (IKI) and TREPA (GCF) projects implemented by Landscape Alliance (CIFOR & ICRAF in action). On *C. eugenioides*, the cherry is the fleshy fruit that contains the seed, so cherry load is a direct measure of fruit and seed output.

Reproductive output across the assessed trees is low, highly variable and weakly related to tree size. The median tree carried an estimated 29 cherries in the whole canopy, 31% of trees carried 15 cherries or fewer, and the coefficient of variation in cherry load exceeded 110%. Tree size explained little of this variation (stem diameter, the single significant structural predictor, accounted for under 10% of the variance; r = 0.31). Trees in the isolated Cyamudongo fragment carried significantly fewer cherries than those in the main Nyungwe block (medians 15 vs 32 cherries; Mann-Whitney p = 0.021), and every observation of sub-optimal on-tree seed quality occurred in the fragment.

Low and asynchronous fruiting, together with the reduced fruit and seed set typical of small, isolated populations, makes large-scale seed collection an unreliable basis for an orchard. The original plan was to collect seed from wild trees and raise a Breeding Seed Orchard (BSO). On the evidence presented here, that plan should be redirected toward a Clonal Seed Orchard (CSO): rather than collecting seed, scions (budwood) are taken from selected wild trees and grafted onto vigorous local Rwandan *Coffea arabica* rootstock. This captures the full genotype of each selected tree, shortens the time to flowering, secures the germplasm of a threatened wild relative *ex situ*, and aligns with Rwanda’s restoration and climate commitments.

## 1. Introduction

### 1.1 Background and rationale

Cultivated Arabica coffee (*Coffea arabica* L.) is an allotetraploid (2n = 4x = 44) that arose from a single, geologically recent hybridisation between two diploid progenitors most closely related to *C. canephora* and *C. eugenioides* (Salojärvi et al., 2024). Because the species originated from very few founding individuals, the cultivated Arabica gene pool carries extremely narrow genetic diversity, which limits the scope for within-species breeding and raises the value of its wild diploid relatives as reservoirs of useful variation (Scalabrin et al., 2020). *C. eugenioides* is the maternal progenitor of Arabica and is one of the comparatively few *Coffea* species native to the Afromontane forests of the Albertine Rift, including Rwanda.

Wild coffee species are under serious threat. A global assessment of all known *Coffea* species concluded that at least 60% are threatened with extinction, that 45% are held in no *ex situ* germplasm collection, and that 28% occur in no protected area, with deforestation and climate change identified as the principal drivers (Davis et al., 2019). For wild Arabica specifically, the application of climate projections shifted the assessment from Least Concern to Endangered, with modelled population reductions of 50% or more by the end of the century (Moat et al., 2019). Conserving and pre-emptively securing crop wild relatives such as *C. eugenioides* is therefore a recognised priority for the long-term resilience of the coffee sector (Davis et al., 2019).

Against this background, a field assessment of wild *C. eugenioides* was carried out in Nyungwe Forest National Park and the Cyamudongo fragment to inform an appropriate seed-source strategy. The original project intention was to establish a Breeding Seed Orchard (BSO): a seedling orchard raised from open-pollinated seed collected from selected wild mother trees, doubling as a progeny test and a seed-production stand (White et al., 2007). Field observations of consistently low seed loads, and of marked asynchrony in fruiting among trees, prompted a re-evaluation of whether seed-based establishment is feasible. This report tests that concern quantitatively and develops an evidence-based alternative.

### 1.2 Study sites

Nyungwe Forest National Park is among the best-preserved montane rainforests in Africa, covering roughly 1,019 km^2^ of the Congo-Nile divide in south-western Rwanda and forming part of the Albertine Rift (Plumptre et al., 2002). Cyamudongo is a small (~19 km^2^) forest fragment lying to the west of the main Nyungwe block, severed from the contiguous forest by decades of agricultural expansion and now surrounded by tea and eucalyptus; despite its isolation it remains administratively part of the national park and retains high biodiversity (Plumptre et al., 2002). The contrast between a large, contiguous forest (Nyungwe) and a small, isolated fragment (Cyamudongo) provides a natural setting in which to examine whether isolation affects the reproductive performance of *C. eugenioides*.

### 1.3 Objectives

The assessment pursued four objectives: (1) to describe the population structure of wild *C. eugenioides* in the two forests through individual-tree measurements of size, fruit production, vigour and health; (2) to quantify reproductive output (cherry load) and phenological status; (3) to map the spatial distribution of trees and their reproductive performance using the georeferenced field data; and (4) to assess, on the basis of this evidence, whether a seed-based Breeding Seed Orchard or a graft-based Clonal Seed Orchard is the more appropriate seed-source strategy.

## 2. Materials and Methods

### 2.1 Study design and sampling

The field assessment targeted mature *C. eugenioides* plus-trees, vigorous, fruit-bearing individuals identifiable as candidate mother trees and therefore as potential seed or scion donors. To avoid over-representing close relatives, selected plus-trees were spaced at least 70 m apart, so that immediate near-neighbours, which are more likely to share recent ancestry and to result from localised, potentially inbred mating, were not sampled together. This minimum-distance criterion reduces spatial genetic autocorrelation among the sampled trees and improves the genetic representativeness of the assessed set. Each qualifying tree was assigned a sequential identifier and recorded as an individual. In total, 59 plus-trees were measured, 42 in the main Nyungwe block and 17 in the Cyamudongo fragment. These trees are a representation of the seed-bearing (fruiting) individuals only; they are not a complete inventory of *C. eugenioides* in either forest, whose total population is larger, particularly in Cyamudongo, where many additional, non-fruiting individuals occur but fell outside the scope of a seed-source assessment.

### 2.2 Data collection instrument

Data were collected digitally with a structured questionnaire built in KoBoToolbox and completed on a mobile device. The instrument was organised as two linked parts: a one-off Seed Source record per site (site name, habitat and vegetation type, land tenure and protection status, population phenology status, and the GPS reference of the seed-source area) and a repeating Individual Tree record completed for every tree. The complete form, every question, its response type, answer options, data-entry hints and skip logic, is reproduced in Annex 1. The principal per-tree variables are summarised in Table 1. On *C. eugenioides*, the cherry is the fleshy fruit enclosing the seed; cherry, fruit and seed therefore refer to the same structure, and the whole-canopy cherry estimate is used throughout as the measure of reproductive output.

**Table 1:**
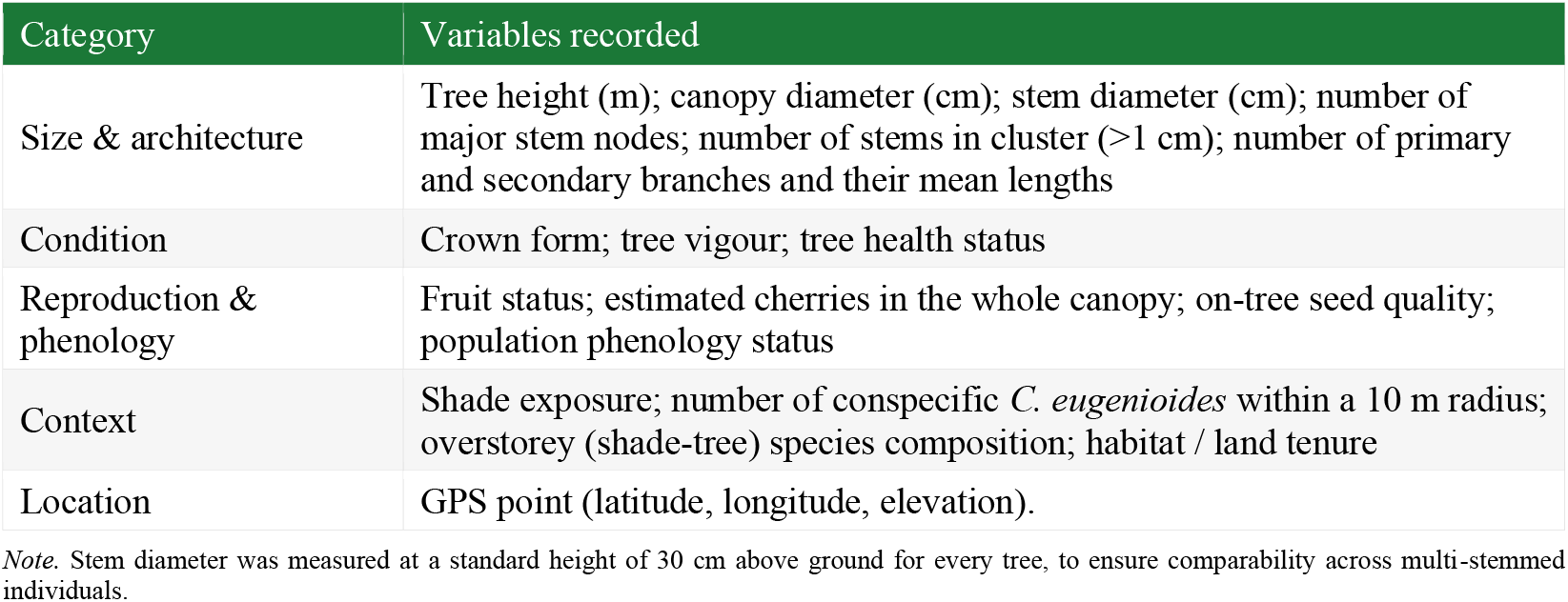
Principal variables recorded for each tree.

| Category | Variables recorded |
| --- | --- |
| Size & architecture | Tree height (m); canopy diameter (cm); stem diameter (cm); number of major stem nodes; number of stems in cluster (>1 cm); number of primary and secondary branches and their mean lengths |
| Condition | Crown form; tree vigour; tree health status |
| Reproduction & phenology | Fruit status; estimated cherries in the whole canopy; on-tree seed quality; population phenology status |
| Context | Shade exposure; number of conspecific *C. eugenoides* within a 10 m radius; overstorey (shade-tree) species composition; habitat / land tenure |
| Location | GPS point (latitude, longitude, elevation). |

Note. Stem diameter was measured at a standard height of 30 cm above ground for every tree, to ensure comparability across multi-stemmed individuals.

### 2.3 Data cleaning

The raw KoBoToolbox export contained 59 genuine tree records followed by a block of stray submission-index numbers in a single column; these non-data rows were removed. Free-text overstorey species names, recorded phonetically in the field, were normalised against accepted botanical binomials (for example, field spellings such as “Strompozia scheffleri” and “crizofilum gorogonzanum” were mapped to *Strombosia scheffleri* and *Chrysophyllum gorungosanum*, respectively).

### 2.4 Site assignment

Each tree was assigned to its forest using the linked Seed Source records: in KoBoToolbox every individual-tree record is nested under the seed-source site at which it was collected, so the recorded site name gives a direct, GPS-independent assignment (the variants “Nyungwe NP”/”Nyungwe” and “Cyamudongo”/”Cyamudongo forest” were merged into Nyungwe and Cyamudongo). This recorded linkage was used in preference to the tree coordinates. On this basis, 42 trees were recorded in Nyungwe and 17 in Cyamudongo.

### 2.5 Statistical analysis

Analyses and figures were produced in R using the *tidyverse, sf, ggplot2, corrplot* and *leaflet* packages. Continuous traits were summarised with means, standard deviations, medians and ranges. Because cherry load was strongly right-skewed, the difference between the two forests was tested with the non-parametric Mann–Whitney *U* test. Pairwise Pearson correlations were computed among structural and reproductive traits, and a multiple linear regression was fitted to identify the strongest predictors of whole-canopy cherry load (predictors: stem diameter, canopy diameter, height, primary and secondary branch counts, conspecific neighbour density, elevation and site). Standardised regression coefficients were used to compare effect sizes on a common scale. Significance was evaluated at *α* = 0.05.

## 3. Results

### 3.1 Population structure and tree condition

The assessed trees were small to medium statured. Tree height averaged 4.0 ± 1.5 m (range 1.7–8.3 m) and stem diameter at 30 cm averaged 3.6 ± 1.6 cm. Canopies were moderately developed (mean canopy diameter ≈ 91 cm), and most individuals were single-stemmed (mean 0.2 stems > 1 cm per cluster). In condition terms the trees were generally sound: 83% were classed as Healthy and the remainder as Moderate, with none in poor health; vigour was split between Medium (53%) and High (47%). Crown form was predominantly Irregular (86%), and trees grew almost exclusively under partial shade (98%). Descriptive statistics are given in Table 2.

**Table 2:**
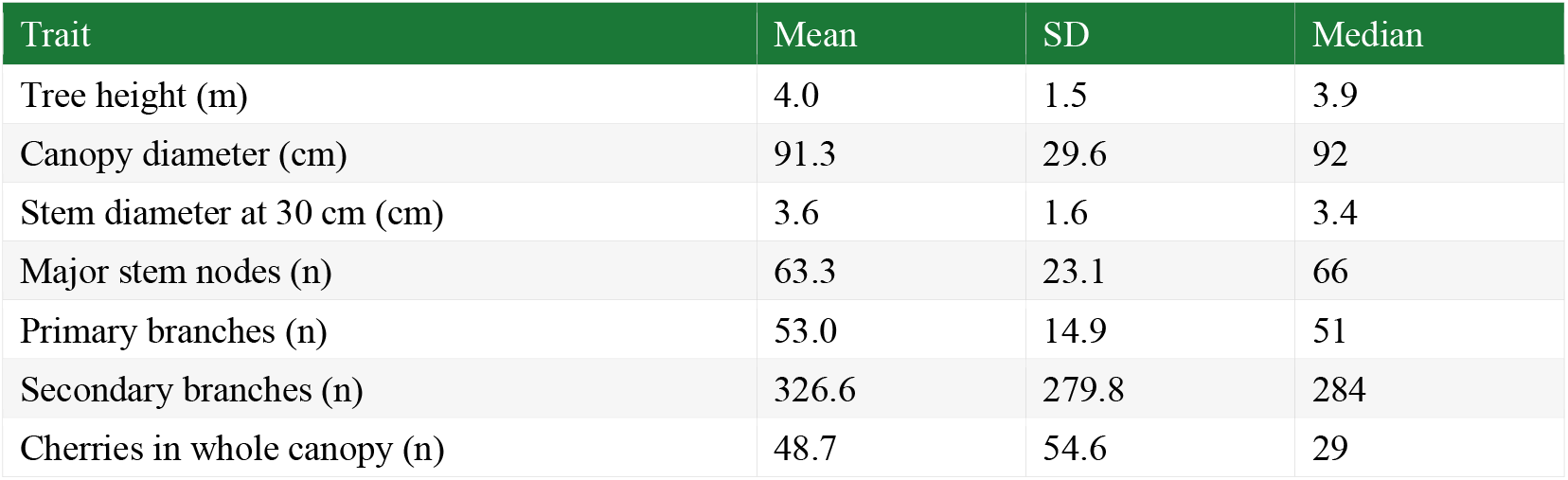
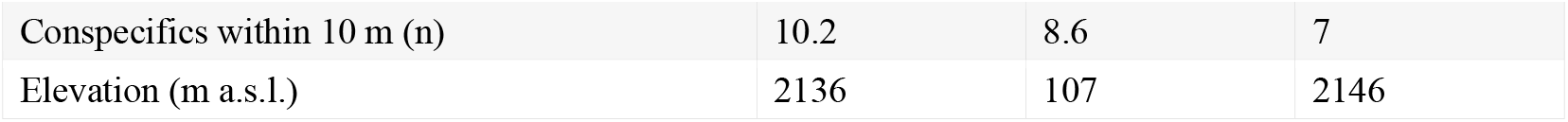
Summary statistics for continuous traits (n = 59 trees).

**Table 3:** Breeding Seed Orchard versus Clonal Seed Orchard for this population.

| Criterion | Breeding Seed Orchard (seedling) | Clonal Seed Orchard (grafted), recommended |
| --- | --- | --- |
| Planting material | Open-pollinated seed from wild mother trees | Scions (budwood) from selected trees |
| Effect of low, asynchronous fruiting | Severe: limits and biases seed collection | Largely avoided, needs only budwood |
| Genetic value captured | Average value via seed | Full genotype of each selected tree |
| Time to flowering / seed | Slow (from seedlings) | Faster (mature scions on established roots) |

The stem-diameter size-class distribution is broadly unimodal, peaking at 2 - 4 cm rather than showing the reverse-J shape (an abundance of small individuals) expected of a vigorously regenerating population (Figure 1). Since the assessment targeted fruit-bearing mother trees rather than all size classes, this pattern is only a preliminary signal of limited recruitment and should be confirmed with dedicated seedling and sapling surveys.

**Figure 1:**
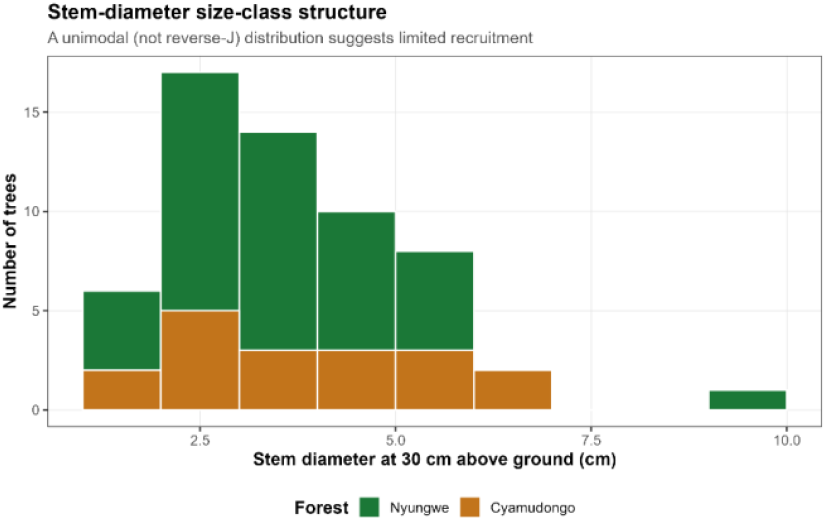
Stem-diameter size-class distribution of the assessed C. eugenioides trees, coloured by forest.

### 3.2 Reproductive output is low and highly variable

Reproductive output was the central concern of the assessment, and the data show it to be low. The median tree carried an estimated 29 cherries in the whole canopy and the mean was 48.7, but the distribution was strongly right-skewed (coefficient of variation > 110%) and dominated by low values: 31% of trees carried 15 cherries or fewer and 53% carried 30 or fewer (Figure 2). Trees field-classed as “Low” fruit status (36% of those assessed) averaged about 12 cherries, with a maximum of 19, while even the most productive trees were a small minority. Across the assessed trees, then, reproductive output is low and clustered near the bottom of the range.

**Figure 2:**
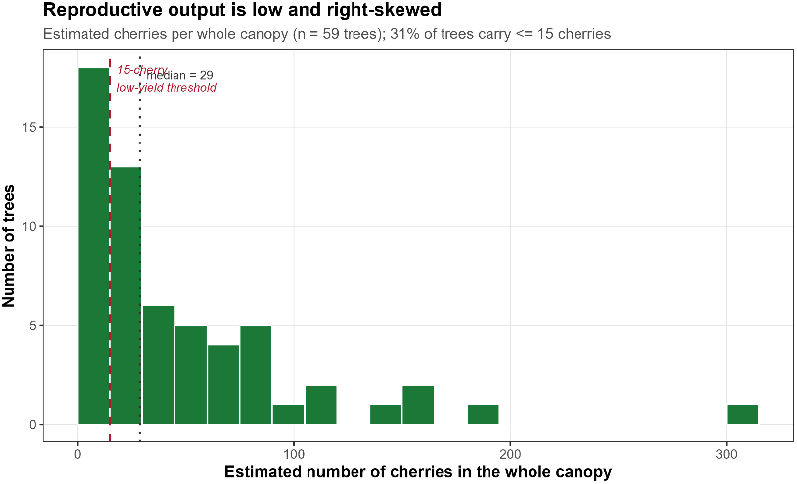
Distribution of estimated whole-canopy cherry load

### 3.3 Production is weakly related to tree size

If low yields simply reflected young or small trees, production should rise steeply with size. It does not. Among all structural traits, stem diameter was the only significant predictor of cherry load in the multiple regression (p = 0.015), yet its simple correlation with cherry load was weak (Pearson r = 0.31), and the full model, including canopy diameter, height, branch counts, conspecific density, elevation and site, explained only about a quarter of the variance (R^2^ ≈ 0.24). Height, branch counts and elevation were all non-significant. The weak stem-diameter relationship (Figure 3) makes the point: larger trees are not reliably more productive, structural size and reproductive output are largely decoupled. Tree size alone is therefore an unreliable guide to which trees will produce abundant seed.

**Figure 3:**
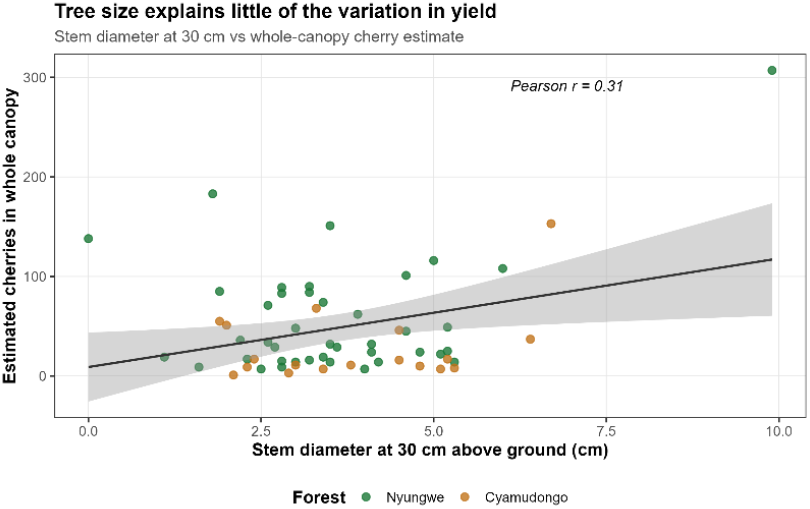
Stem diameter at 30 cm versus whole-canopy cherry load, with linear fit.

### 3.4 The isolated fragment produces less, with poorer seed quality

Trees in the isolated Cyamudongo fragment carried significantly fewer cherries than those in the contiguous Nyungwe block (medians 15 vs 32 cherries; means 29.5 vs 56.5; Mann-Whitney U = 495, p = 0.021; Figure 4). The contrast in seed quality was sharper still: every sub-optimal rating occurred in Cyamudongo, 24% of the total assessed trees there were below ‘Good’ (18%: Average and 6%: Poor), whereas all Nyungwe trees were rated ‘Good’ (Figure 5). These results are consistent with the reduced seed set typical of small, isolated plant populations.

**Figure 4:**
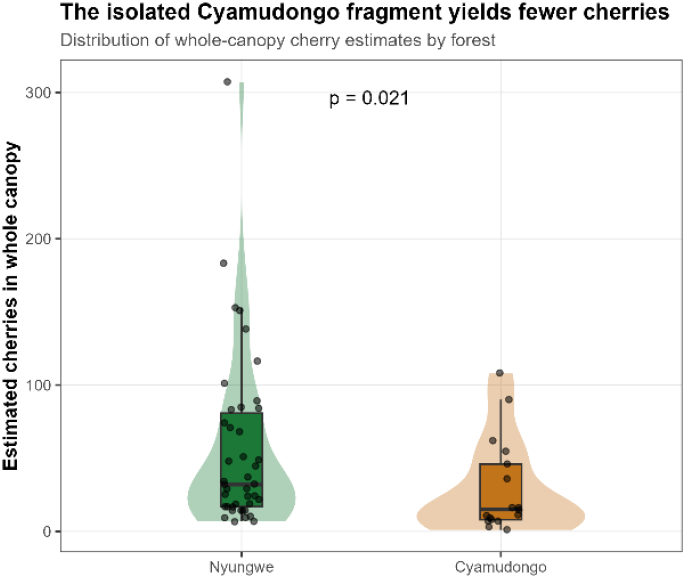
Whole-canopy cherry load by forest.

**Figure 5:**
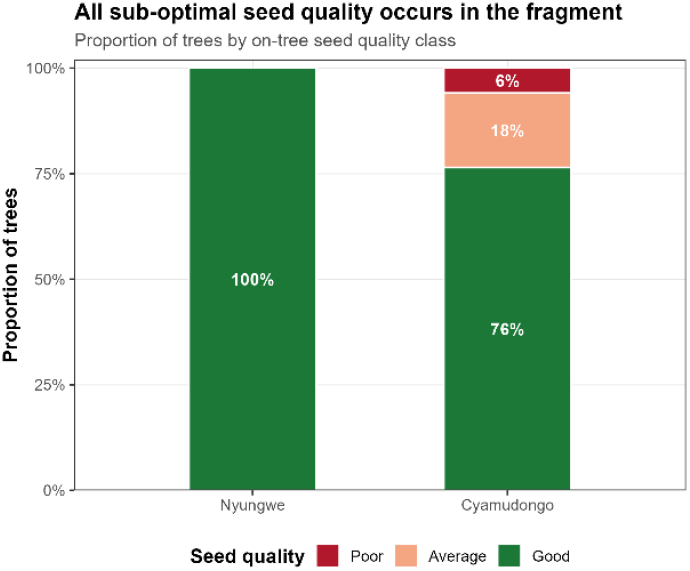
On-tree seed-quality composition by forest.

### 3.5 Spatial distribution

The georeferenced records resolve into the two expected clusters, the main Nyungwe block to the east and the Cyamudongo fragment to the west, spanning an elevation gradient of roughly 1,860 - 2,350 m a.s.l. Mapping per-tree cherry load (Figure 6) makes the spatial pattern explicit: the higher-yielding trees are concentrated in Nyungwe, while the fragment is dominated by low-yielding trees; cherry load shows no systematic trend with elevation.

**Figure 6:**
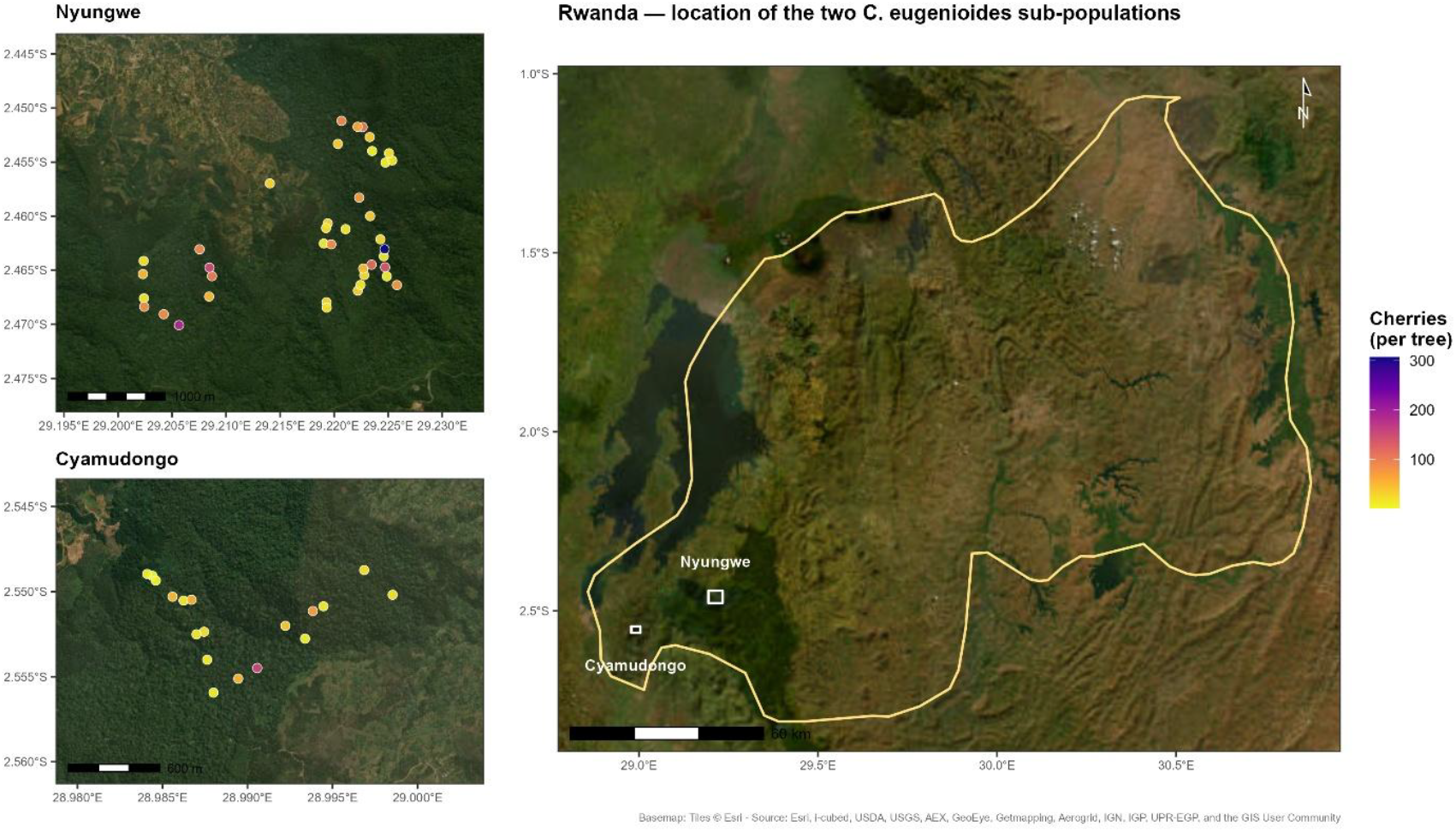
Spatial distribution of assessed trees. (Right) Rwanda overview with national boundary and two subpopulations boxed; (left) Zoomed views of Nyungwe and Cyamudongo Forests.

### 3.6 Habitat context and overstorey associates

Trees occurred overwhelmingly under partial shade within closed montane forest. Conspecific density was modest and variable (mean 10.2, median 7 other *C. eugenioides* within a 10 m radius) and was somewhat higher in Nyungwe than in Cyamudongo. The most frequently recorded overstorey associates were *Strombosia scheffleri, Chrysophyllum gorungosanum, Carapa grandiflora, Newtonia buchananii* and *Parinari excelsa* (Figure 7), a characteristic Albertine Rift montane assemblage that defines the light and microclimate in which *C. eugenioides* grows naturally and which any orchard or enrichment planting should seek to emulate.

**Figure 7:**
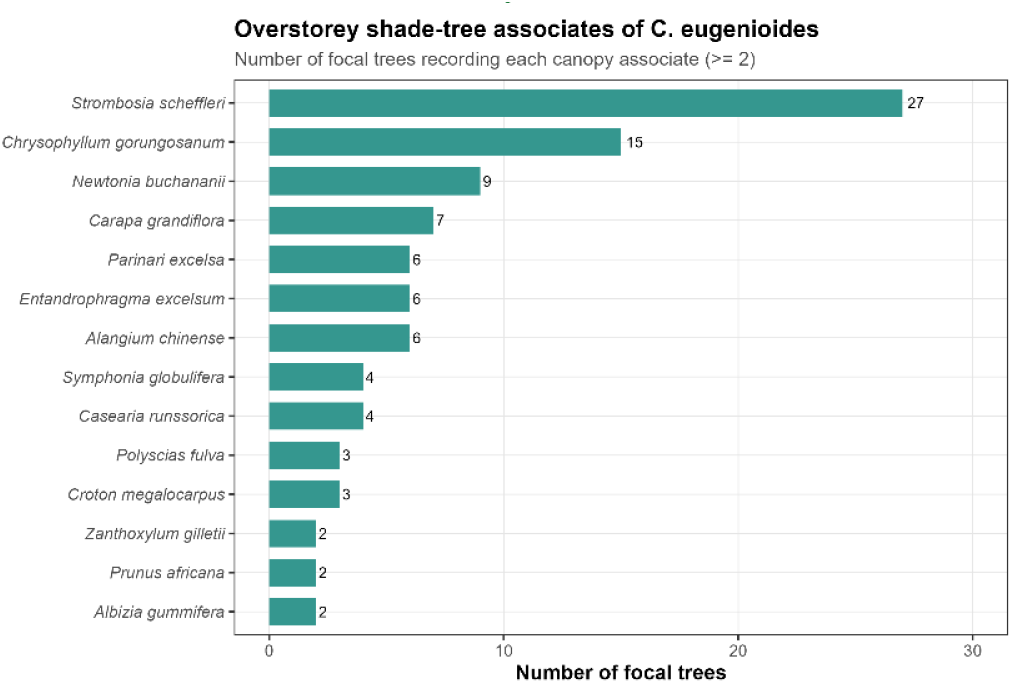
Most frequently recorded overstorey shade-tree associates of C. eugenioides (species recorded for ≥ 2 trees).

**Figure 8:**
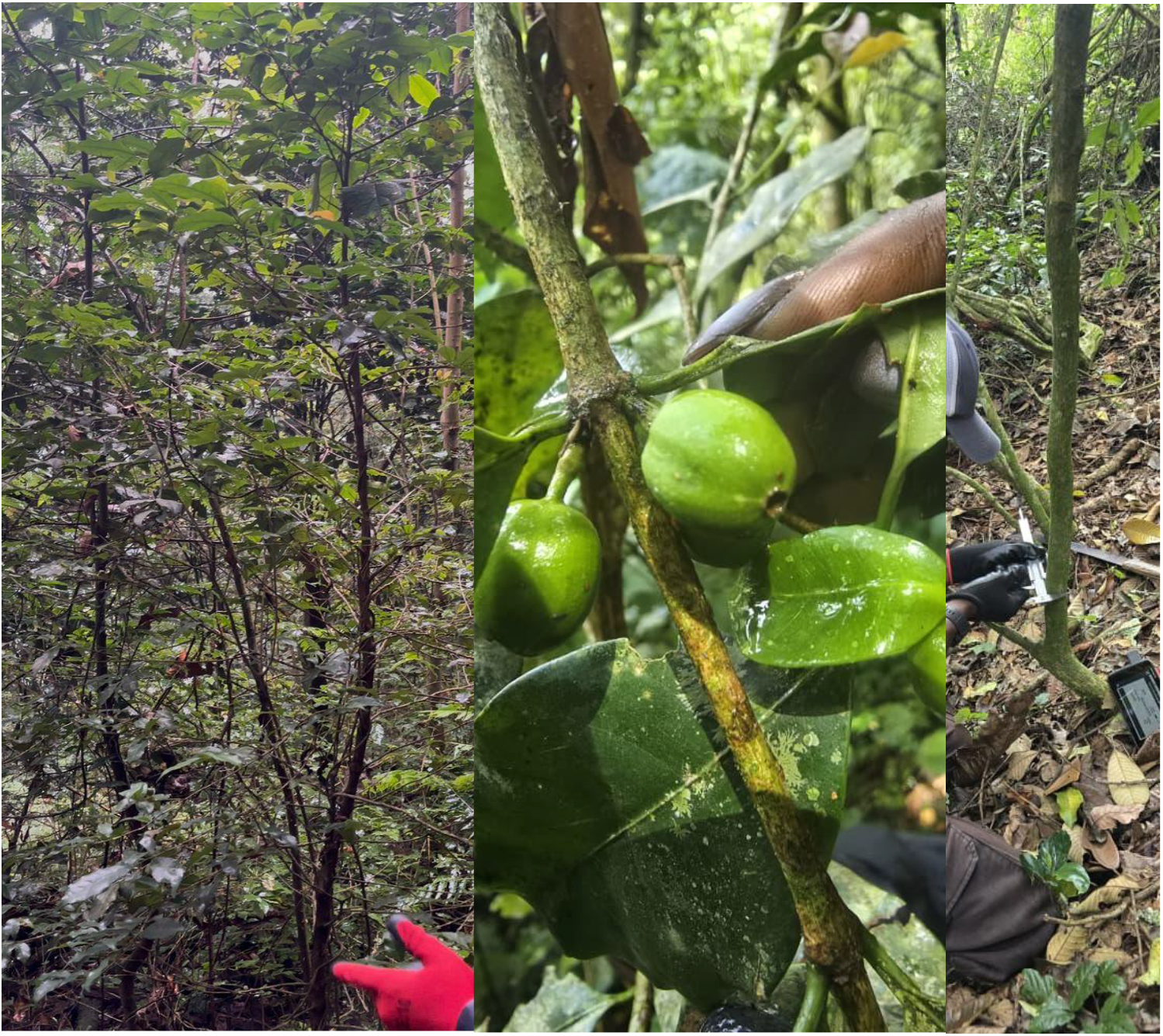
C. eugenioides tree (left), cherry count (middle), stem diameter and geolocation recording (right).

## 4. Discussion

The low cherry loads reported here are best judged against cultivated coffee. Under good management an Arabica bush is a heavy bearer, with mature plants carrying many hundreds to several thousand fruits per year (DaMatta et al., 2007), whereas the wild *C. eugenioides* mother trees assessed here carried a median of only 29 whole-canopy cherries (maximum 307). Wild, shaded, undomesticated understorey trees are admittedly not directly comparable to intensively managed bushes; nonetheless the order-of-magnitude difference confirms that the standing crop on any single wild tree is far too small to support reliable bulk seed collection. This modest productivity does not diminish the value of the species, if anything it sharpens it. *C. eugenioides* is one of the two diploid progenitors of Arabica and a reservoir of variation absent from the narrow cultivated gene pool (Scalabrin et al., 2020; Salojärvi et al., 2024). It accumulates only about half the caffeine of Arabica (roughly 0.5% of dry matter) together with comparatively low chlorogenic acids, the compounds responsible for much of coffee’s bitterness, giving a milder cup and making it an attractive parent for breeding low-caffeine, higher-quality coffee (Campa et al., 2005; Salojärvi et al., 2024). As a climate-threatened crop wild relative, it is also a potential source of disease-resistance and climate-resilience traits for a sector under mounting pressure (Davis et al., 2019; Aerts et al., 2013). The orchard’s purpose is therefore to capture and secure these high-value genotypes, not merely to conserve an endangered plant, but to keep a uniquely valuable genetic resource available for future coffee improvement.

### 4.1 Why is production so low and variable?

Three lines of evidence converge on the conclusion that the limitation on these trees is reproductive rather than simply a matter of immature or small individuals. First, yields are low in absolute terms and concentrated near the bottom of the range. Second, yield is only weakly explained by tree size; the strongest size predictor, stem diameter, accounts for under 10% of the variance. Third, the lowest yields and the only sub-optimal seed-quality ratings occur in the small, isolated Cyamudongo fragment. Together, these patterns point toward the reduced pollination and fruit set characteristic of small, fragmented plant populations.

A large body of work shows that habitat fragmentation reduces the reproductive success of animal-pollinated plants. A meta-analysis spanning many species found a strong, consistently negative effect of fragmentation on both pollination and reproduction, with self-incompatible and specialised species most affected (Aguilar et al., 2006). Small, isolated remnants typically support fewer and less diverse pollinators, reducing pollen deposition and depressing fruit and seed set and quality, and they show measurable shifts in tree reproductive traits relative to continuous forest (Girão et al., 2007; Hadley & Betts, 2012). For coffee specifically, fruit set in *Coffea arabica*, although the species is capable of self-fertilisation, increases markedly with the diversity and abundance of pollinating bees; recent experimental work records about 9% higher fruit set under bee pollination, alongside quality gains (Klein et al., 2003; Aristizábal et al., 2025).

The lower cherry load and poorer seed quality in Cyamudongo are consistent with this mechanism: a small, isolated population with a reduced or unstable pollinator service.

Asynchronous flowering compounds the problem. Field observations indicated that trees do not all flower and fruit at the same time, therefore, a single seed-collection visit encounters only a fraction of trees bearing mature fruit. Asynchrony of this kind reduces the pollen available for outcrossing and lowers the effective number of trees contributing to any one seed lot (Aguilar et al., 2006; DaMatta et al., 2007). For an operational seed-collection programme, this means that the realised seed harvest per visit is far smaller, and far less genetically representative, than the standing number of trees would suggest.

### 4.2 Implications for the seed-source strategy

A Breeding (or seedling) Seed Orchard depends on collecting adequate, approximately balanced quantities of seed from many selected mother trees, to ensure that the resulting orchard captures broad genetic diversity and can double as a progeny test (White et al., 2007). The conditions documented here undermine that model in three ways. (i) Quantity: with a median of 29 cherries per tree and a third of trees below 15, the seed available per mother tree is low. (ii) Reliability and representativeness: asynchronous fruiting makes balanced, simultaneous collection across the selected trees impractical, biasing any seed lot toward whichever few trees happen to be fruiting. (iii) Seed quality: sub-optimal on-tree seed quality in the fragment, together with the elevated inbreeding expected in small populations, raises the risk that collected seed yields weak or genetically narrow progeny (Aguilar et al., 2006; Girão et al., 2007). These are precisely the circumstances under which a clonal, rather than seedling, founding strategy is preferred.

### 4.3 The case for a Clonal Seed Orchard

A Clonal Seed Orchard (CSO) is established by vegetatively propagating selected plus-trees, most commonly by grafting scions onto rootstock, and arranging the resulting ramets to promote wide outcrossing and heavy seed production (White et al., 2007). For a species in the situation of these *C. eugenioides* trees, the clonal route has decisive advantages:

✓ It bypasses the seed bottleneck. Establishing the orchard does not require large, balanced seed lots from a low-yielding, asynchronously fruiting wild population; it requires only scions (budwood), which can be taken whenever suitable shoots are available.
✓ It captures the entire selected genotype. Grafting reproduces the full genotype of each plus-tree, rather than only the average value transmitted through open-pollinated seed, and brings superior genotypes into production sooner (White et al., 2007).
✓ It shortens time to flowering. Scions taken from mature, already-reproductive trees and grafted onto established rootstock flower far sooner than seedlings, accelerating both seed production and any future controlled crossing.
✓ It secures threatened germplasm ex situ. Because wild Coffea is highly threatened (Davis et al., 2019), a grafted living collection provides an immediate, individually documented ex-situ safeguard of identified wild genotypes, a recognised complementary conservation strategy for the genus (Aerts et al., 2013; Krishnan et al., 2013).

### 4.4 Grafting *C. eugenioides* onto local Coffea arabica rootstock

Grafting is an operationally routine technique in coffee. Interspecific grafting within *Coffea*: most commonly Arabica scions on *C. canephora* (robusta) rootstock, is used to confer nematode resistance, drought tolerance and a more vigorous root system (Bertrand et al., 2001; Koutouleas et al., 2023). The proposal here is the converse pairing: placing *C. eugenioides* scions onto locally adapted Rwandan *C. arabica* rootstock. Several points support its feasibility. *C. eugenioides* is one of the two diploid progenitors of Arabica and is genomically very close to the Arabica E-subgenome (Salojärvi et al., 2024), so graft compatibility is biologically plausible. Locally selected Arabica is widely available in Rwanda, is adapted to local soils and climate, and provides a robust root system.

Interspecific rootstock - scion combinations can modify vigour and yield relative to non-grafted plants, and outcomes vary with the specific genotypes used (Bertrand et al., 2001; Koutouleas et al., 2023). Grafting onto Arabica also does not by itself broaden the genetic base, that depends on how many distinct wild trees are sampled as scion donors, therefore, capturing diversity remains a design requirement.

### 4.5 Limitations

Several limitations temper interpretation. Cherry load was recorded as a rapid whole-canopy field estimate rather than an exhaustive count, and is therefore approximate; nonetheless the gradient from low to high is informative and the site contrast is statistically reliable. The assessed trees are fruit-bearing mother-tree candidates rather than a random sample of all individuals, and the two forests were unequally represented, so the figures describe the seed-source pool rather than the whole population. Flowering asynchrony was observed qualitatively rather than quantified through repeated monitoring, and the lower production in Cyamudongo, while consistent with isolation effects, is correlational and could be influenced by elevation, soil or microclimate. Pollinator communities were not surveyed, and the inferred pollination limitation, although well supported by the literature, was not measured directly.

## 5. Conclusions and Recommendations

### 5.1 Conclusions

The assessed mother trees are structurally sound, small to medium statured, predominantly healthy and of medium-to-high vigour, but the size-class profile gives a preliminary signal of limited recruitment among fruit-bearing trees. Cherry load is low, highly variable and only weakly related to tree size, and fruiting is asynchronous across individuals. The trees form two clear clusters along a 1,860 to 2,350 m elevation gradient, with the higher-yielding trees concentrated in Nyungwe and the isolated Cyamudongo fragment producing significantly less and showing every case of sub-optimal seed quality. These conditions make a seed-based Breeding Seed Orchard slow, unreliable and genetically risky, whereas a graft-based Clonal Seed Orchard avoids the seed bottleneck, captures the full genotype of selected trees and secures the germplasm *ex situ*. The evidence therefore supports a Clonal Seed Orchard, established by grafting selected wild *C. eugenioides* scions onto locally adapted Rwandan *C. arabica* rootstock, as the more feasible strategy.

### 5.2 Recommendation 1

Re-orient the orchard objective from a seedling BSO to a grafted CSO. Select donor trees on the basis of demonstrated fruiting, health and vigour rather than size alone, and collect scions (budwood) for grafting onto established local *C. arabica* rootstock in a managed nursery. Field-plant the resulting ramets in a wide-spaced layout that keeps ramets of the same clone apart, to promote outcrossing and heavy seed production (White et al., 2007).

### 5.3 Recommendation 2

Because cultivated Arabica and its wild relatives already have narrow diversity (Scalabrin et al., 2020), the orchard should be established on as many distinct, unrelated wild trees as can be sourced, with donors drawn from both forests and across the elevation range, and each donor tree permanently georeferenced. Sampling broadly at the outset is the simplest safeguard against inbreeding and is consistent with established guidance on sourcing germplasm for restoration and conservation (Thomas et al., 2014; FAO, 2014).

### 5.4 Alignment with Rwanda’s priorities

The strategy aligns directly with national and project commitments. Rwanda’s restoration agenda under the Bonn Challenge and the African Forest Landscape Restoration Initiative, emphasises well-adapted native species, and the country’s Nationally Determined Contribution prioritises ecosystem restoration and climate-resilient land use (Republic of Rwanda, 2020). RTRP-Seed exists precisely to raise the availability and quality of native tree germplasm, while TREPA restores degraded landscapes with native and agroforestry species; both are implemented by Landscape Alliance (CIFOR & ICRAF in action). A documented, grafted clonal orchard of a threatened native crop wild relative is a concrete contribution to the native-species seed systems these projects are designed to build.

## Acknowledgements

This assessment was undertaken in May 2026, under the Right Tree in the Right Place for the Right Purpose Project (RTRP-Seed), funded by the International Climate Initiative (IKI), and the Transforming Eastern Province through Adaptation (TREPA) project, led by The International Union for Conservation of Nature (IUCN), funded by the Green Climate Fund (GCF). Both projects are implemented by the Landscape Alliance (CIFOR & ICRAF in action). The support of these projects and of the Rwanda Forest Authority (RFA), and the management of Nyungwe National Park for facilitating access to the field sites, is gratefully acknowledged.

Sincere thanks are extended to the reviewers listed below, whose careful reading and technical input substantially improved this report.

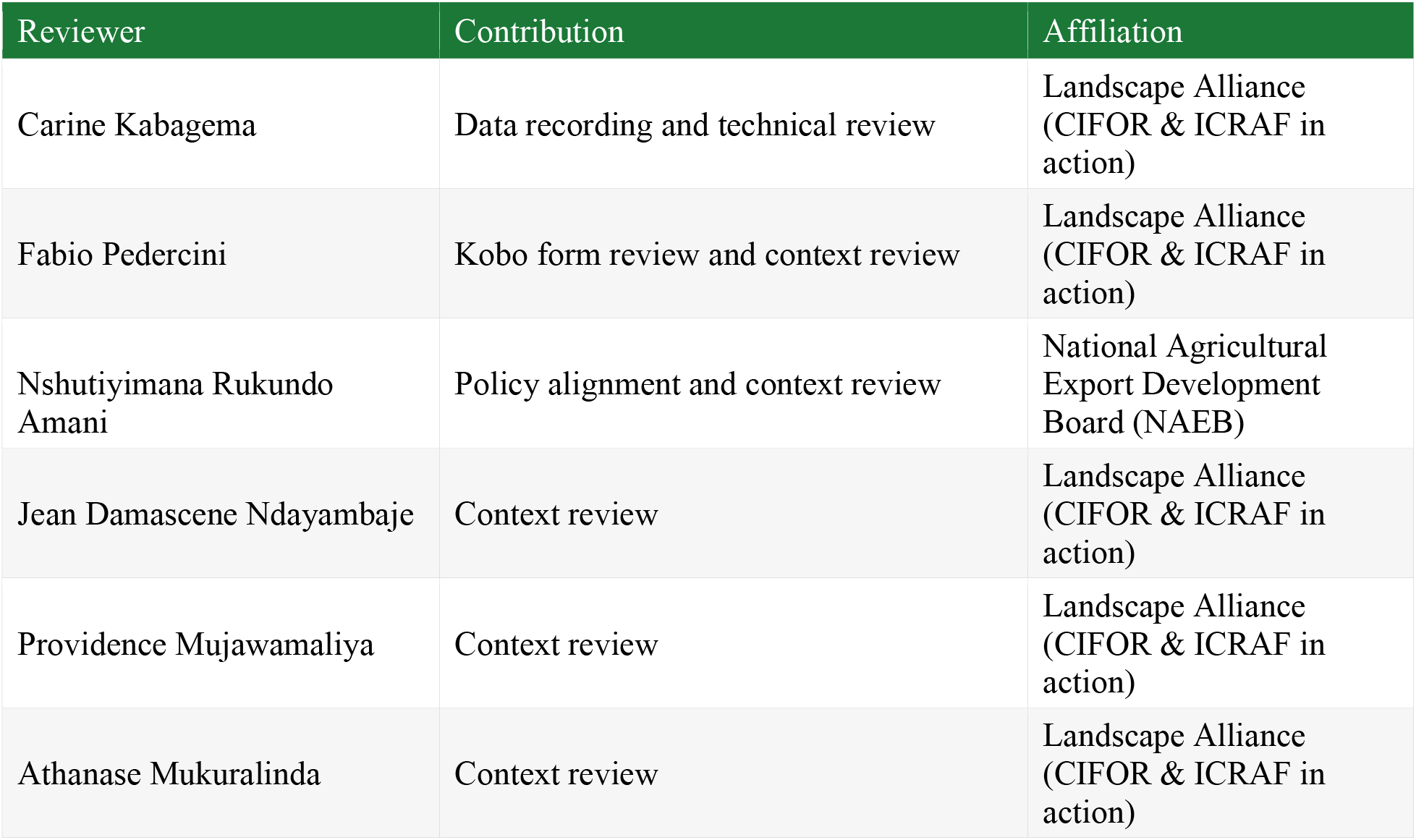

| Reviewer | Contribution | Affiliation |
| --- | --- | --- |
| Carine Kabagema | Data recording and technical review | Landscape Alliance (CIFOR & ICRAF in action) |
| Fabio Pedercini | Kobo form review and context review | Landscape Alliance (CIFOR & ICRAF in action) |
| Nshutiyimana Rukundo Amani | Policy alignment and context review | National Agricultural Export Development Board (NAEB) |
| Jean Damascene Ndayambaje | Context review | Landscape Alliance (CIFOR & ICRAF in action) |
| Providence Mujawamaliya | Context review | Landscape Alliance (CIFOR & ICRAF in action) |
| Athanase Mukuralinda | Context review | Landscape Alliance (CIFOR & ICRAF in action) |

## Annexes

### Annex 1: Field data-collection form (KoBoToolbox)

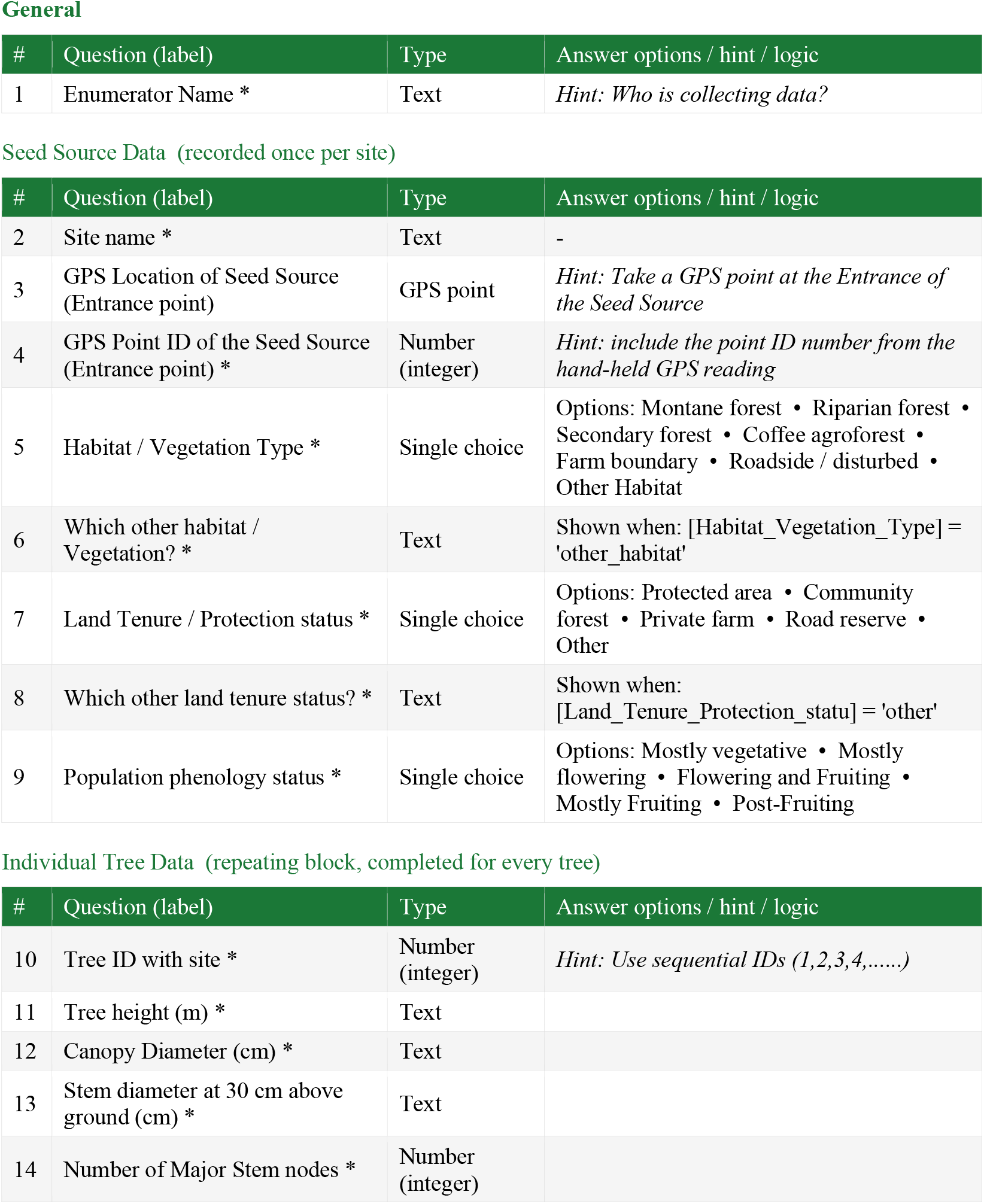
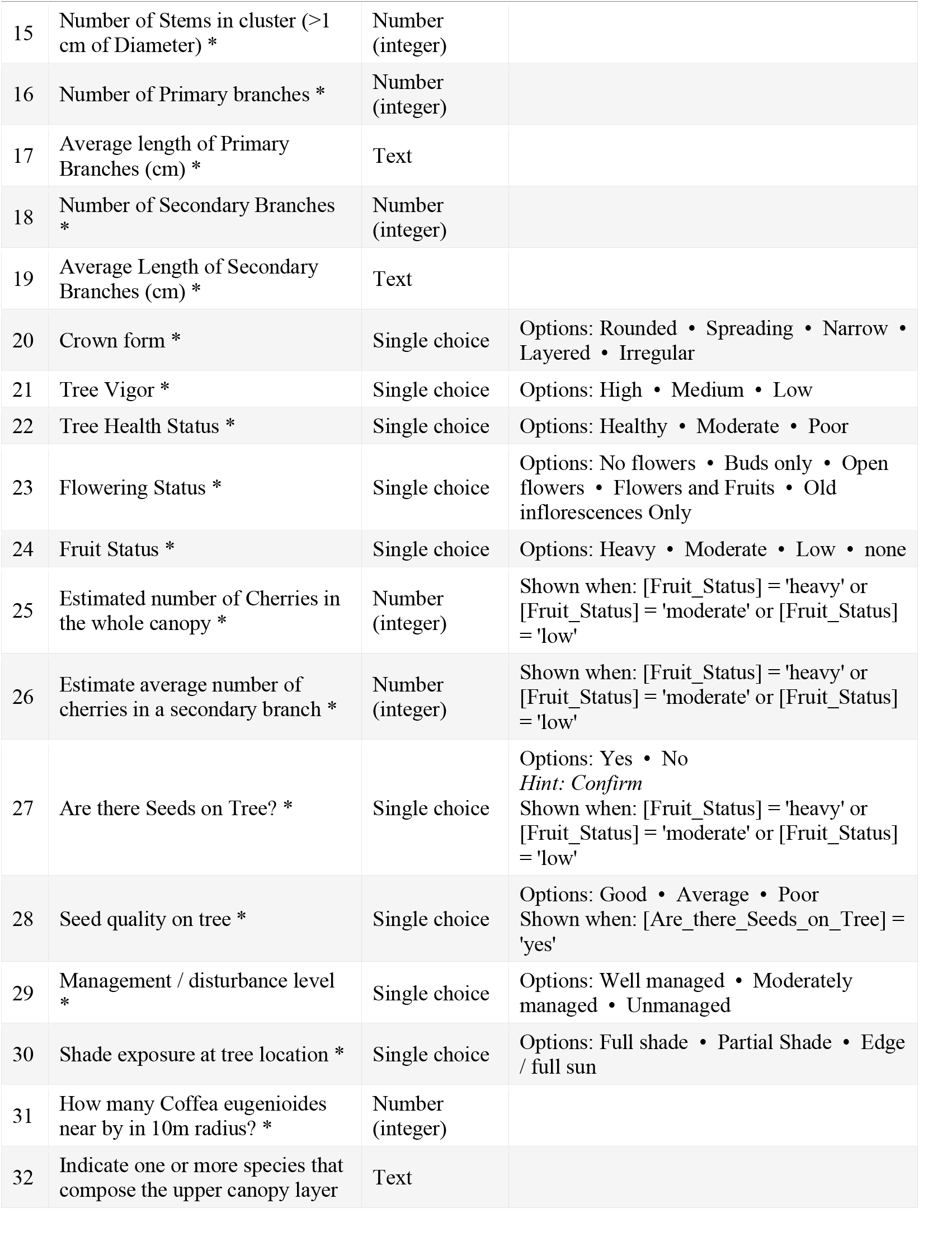
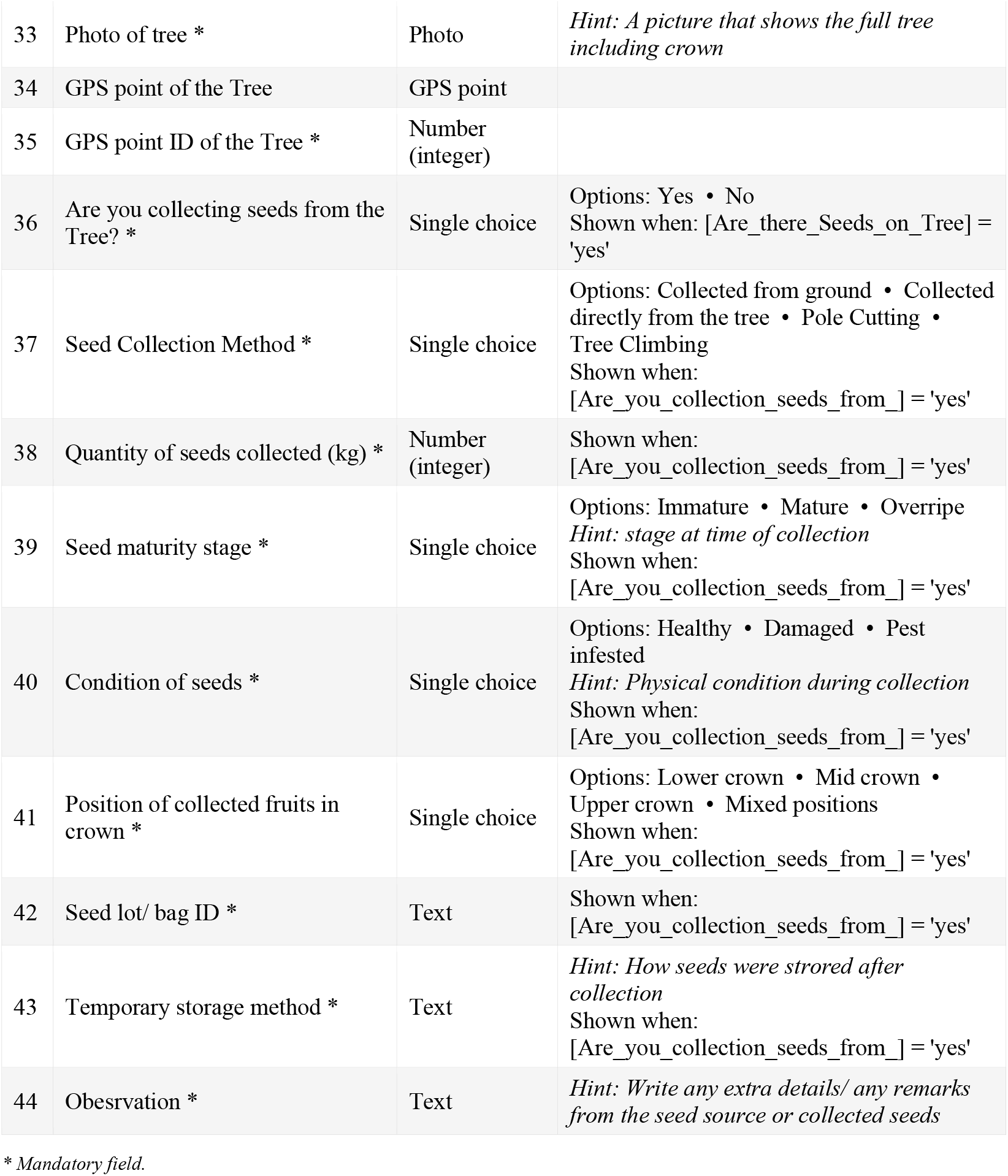

General
| # | Question (label) | Type | Answer options / hint / logic |
| --- | --- | --- | --- |
| 1 | Enumerator Name * | Text | *Hint: Who is collecting data?* |

Seed Source Data (recorded once per site)
| # | Question (label) | Type | Answer options / hint / logic |
| --- | --- | --- | --- |
| 2 | Site name * | Text | - |
| 3 | GPS Location of Seed Source (Entrance point) | GPS point | *Hint: Take a GPS point at the Entrance of the Seed Source* |
| 4 | GPS Point ID of the Seed Source (Entrance point) * | Number (integer) | *Hint: include the point ID number from the hand-held GPS reading* |
| 5 | Habitat / Vegetation Type * | Single choice | Options: Montane forest • Riparian forest • Secondary forest • Coffee agroforest • Farm boundary • Roadside / disturbed • Other Habitat |
| 6 | Which other habitat / Vegetation? * | Text | Shown when: [Habitat_Vegetation_Type] = 'other_habitat' |
| 7 | Land Tenure / Protection status * | Single choice | Options: Protected area • Community forest • Private farm • Road reserve • Other |
| 8 | Which other land tenure status? * | Text | Shown when: [Land_Tenure_Protection_statu] = 'other' |
| 9 | Population phenology status * | Single choice | Options: Mostly vegetative • Mostly flowering • Flowering and Fruiting • Mostly Fruiting • Post-Fruiting |

Individual Tree Data (repeating block, completed for every tree)
| # | Question (label) | Type | Answer options / hint / logic |
| --- | --- | --- | --- |
| 10 | Tree ID with site * | Number (integer) | *Hint: Use sequential IDs (1,2,3,4,.....)* |
| 11 | Tree height (m) * | Text |  |
| 12 | Canopy Diameter (cm) * | Text |  |
| 13 | Stem diameter at 30 cm above ground (cm) * | Text |  |
| 14 | Number of Major Stem nodes * | Number (integer) |  |
| 15 | Number of Stems in cluster (>1 cm of Diameter) * | Number (integer) |  |
| 16 | Number of Primary branches * | Number (integer) |  |
| 17 | Average length of Primary Branches (cm) * | Text |  |
| 18 | Number of Secondary Branches * | Number (integer) |  |
| 19 | Average Length of Secondary Branches (cm) * | Text |  |
| 20 | Crown form * | Single choice | Options: Rounded • Spreading • Narrow • Layered • Irregular |
| 21 | Tree Vigor * | Single choice | Options: High • Medium • Low |
| 22 | Tree Health Status * | Single choice | Options: Healthy • Moderate • Poor |
| 23 | Flowering Status * | Single choice | Options: No flowers • Buds only • Open flowers • Flowers and Fruits • Old inflorescences Only |
| 24 | Fruit Status * | Single choice | Options: Heavy • Moderate • Low • none |
| 25 | Estimated number of Cherries in the whole canopy * | Number (integer) | Shown when: [Fruit_Status] = 'heavy' or [Fruit_Status] = 'moderate' or [Fruit_Status] = 'low' |
| 26 | Estimate average number of cherries in a secondary branch * | Number (integer) | Shown when: [Fruit_Status] = 'heavy' or [Fruit_Status] = 'moderate' or [Fruit_Status] = 'low' |
| 27 | Are there Seeds on Tree? * | Single choice | Options: Yes • No<br>*Hint: Confirm*<br>Shown when: [Fruit_Status] = 'heavy' or [Fruit_Status] = 'moderate' or [Fruit_Status] = 'low' |
| 28 | Seed quality on tree * | Single choice | Options: Good • Average • Poor<br>Shown when: [Are_there_Seeds_on_Tree] = 'yes' |
| 29 | Management / disturbance level * | Single choice | Options: Well managed • Moderately managed • Unmanaged |
| 30 | Shade exposure at tree location * | Single choice | Options: Full shade • Partial Shade • Edge / full sun |
| 31 | How many Coffea eugenioides near by in 10m radius? * | Number (integer) |  |
| 32 | Indicate one or more species that compose the upper canopy layer | Text |  |
| 33 | Photo of tree * | Photo | *Hint: A picture that shows the full tree including crown* |
| 34 | GPS point of the Tree | GPS point |  |
| 35 | GPS point ID of the Tree * | Number (integer) |  |
| 36 | Are you collecting seeds from the Tree? * | Single choice | Options: Yes • No<br>Shown when: [Are_there_Seeds_on_Tree] = 'yes' |
| 37 | Seed Collection Method * | Single choice | Options: Collected from ground • Collected directly from the tree • Pole Cutting • Tree Climbing<br>Shown when: [Are_you_collection_seeds_from_] = 'yes' |
| 38 | Quantity of seeds collected (kg) * | Number (integer) | Shown when: [Are_you_collection_seeds_from_] = 'yes' |
| 39 | Seed maturity stage * | Single choice | Options: Immature • Mature • Overripe<br>*Hint: stage at time of collection*<br>Shown when: [Are_you_collection_seeds_from_] = 'yes' |
| 40 | Condition of seeds * | Single choice | Options: Healthy • Damaged • Pest infested<br>*Hint: Physical condition during collection*<br>Shown when: [Are_you_collection_seeds_from_] = 'yes' |
| 41 | Position of collected fruits in crown * | Single choice | Options: Lower crown • Mid crown • Upper crown • Mixed positions<br>Shown when: [Are_you_collection_seeds_from_] = 'yes' |
| 42 | Seed lot/ bag ID * | Text | Shown when: [Are_you_collection_seeds_from_] = 'yes' |
| 43 | Temporary storage method * | Text | *Hint: How seeds were stored after collection*<br>Shown when: [Are_you_collection_seeds_from_] = 'yes' |
| 44 | Obesrvation * | Text | *Hint: Write any extra details/ any remarks from the seed source or collected seeds* |

\* Mandatory field.

### Annex 2: Field photographs

